# Identification of molecular specificity between reprogrammed cardiomyocytes and normal cardiomyocytes by bioinformatics

**DOI:** 10.1101/2020.04.18.045955

**Authors:** Mengyao Liu, yue Zhang

## Abstract

The adult human heart lacks an effective endogenous repair mechanism and cannot fully restore heart function after injury. We obtained two microarray data sets GSE99814 and GSE22292 from the NCBI GEO database, which include 5 groups of directly reprogrammed cardiomyocyte-like cells (iCMs) and 5 groups of normal mouse cardiomyocytes (CM). And through the GEO2R tool and Venn diagram software to screen the differentially expressed genes (DEG) between iCM and CM. Next, we performed functional enrichment analysis on these DEGs. The protein-protein interaction network (PPI) was constructed by STRING and Cytoscape for module analysis. We have selected a total of 243 DEGs consistently expressed genes in the two data sets, of which 127 up-regulated genes function and pathway enrichment mainly concentrated on biological processes such as innate immune response, inflammatory response, immune system process, positive regulation of apoptosis process and complement and coagulation cascade, while the 116 down-regulated genes are mainly enriched in cell cycle, cell division, cardiac development, myocardial contraction and myocardial cell development and signaling pathways such as cell cycle and adrenergic signaling in cardiomyocytes. Then in the PPI network, we found 27 central genes when analyzing these 243 DEGs by the Molecular Complex Detection (MCODE) plug-in. Finally, we reanalyzed these 27 central genes through DAVID and found that 6 genes (CCNB1, CCNB2, BUB1, TTK, CDC25C, CCNA2) significantly enriched the cell cycle signaling pathway (*p* <1.20E-07). Therefore, through integrated bioinformatics methods, we found that compary with iCMs and CMs, iCMs were induced by direct reprogramming of neonatal mouse cardiomyocytes, the DEGs were mainly enriched in immune-related processes and myocardial contraction and cell cycle signaling pathways (*p* < 0.05).

## Introduction

Heart failure (HF) remains the leading cause of human death worldwide. Mammal adult hearts have little ability to regenerate after injury. For severe heart failure patients, the effect of drug treatment is very limited. Therefore, functional cardiomyocytes (CMs) or CM-like cells produced by replacement therapy will provide an alternative strategy for repairing damaged hearts. Direct in vivo cardiac reprogramming is to inject the reprogramming mixture directly into the injured heart without the need for cell transplantation but can be regenerated in situ. This is a technique with great potential to transform damaged myocardial tissue into functional cardiomyocytes. One of them is inducing cardiac differentiation by pluripotent stem cells (PSC), such as inducing embryonic stem cells (ESC) and induced pluripotent stem cells (iPSC) into CM^[1]^, and the other is directly reprogramming cardiac fibroblast (CF) to CM. Many studies had shown that humans can directly induce mouse embryonic fibroblasts (MEF), tail tip fibroblast (TFF) or CF to reprogram into iCM by using transcription factors GMT^[2,8]^, microRNA-133^[3]^ and chemical mixtures^[4]^. Recently, in vivo studies have found that Sendai virus (SeV) can effectively and rapidly express cardiac reprogramming factors, and rapidly reprogram mouse and human fibroblasts to integration-free iCM. In addition, SeV-GMT can be reversed than GMT Recording virus produces more than 100 times higher pulsating cardiomyocytes^[5]^. Similarly, there is also a evidence that direct cardiac reprogramming can reduce scar tissue from damaged myocardium and generate new cardiomyocytes to restore cardiac function, so direct cardiac reprogramming can be used as an effective strategy for cardiac regeneration. However, there are still some problems before the direct reprogramming method is used in clinical treatment, including the low conversion rate of iCMs, the need for reprogramming in the case of acute injury^[6]^, and the tumorigenicity caused by in vivo reprogramming induction. Therefore, in this study, bioinformatics methods were used to compare and analyze directly reprogrammed cardiomyocyte-like cells induced by mouse myocardial fibroblasts (CF) from the same source. Compared with normal cardiomyocytes, we found the up-regulated DEGs in iCM are significantly enriched in biological processes related to immunology, and the down-regulated DEGs are significantly enriched in myocardial contraction and cell cycle signaling pathways, which may be the key to reprogramming fibroblasts into cardiomyocytes.

## Methods

### Microarray data information

NCBI-GEO is a free public database of microarray / gene profiles. In order to ensure the same genetic background, we obtained two Gene expression data of GSE99814 and GSE22292, which were reprogrammed directly by cardiac fibroblasts (CFs) of α – MHC-GFP transgenic newborn mice [1], into cardiomyocytes like cells (iCMs). The chip data of GSE99814 comes from the platform of GPL13912 (Agilent-028005 Sure Print G3 Mouse GE 8×60K Microarray (Feature Number version)), the data chip of GSE22292 comes from the platform of GPL6246 ([MoGene-1_0-st] Affymetrix Mouse Gene 1.0 ST Array [transcript (gene) version]), including five groups of myocardial like cells induced by GMT (GATA4, MEF2C, TBX5) and five groups of mouse myocardial cells. They include 5 groups of GMT(Gata4, Mef2c, Tbx5) induced cardiomyocytes and 5 groups of mouse cardiomyocytes.

### DEGs data processing

The differentially expressed genes (DGEs) between the iCM samples and the mouse cardiomyocyte samples was obtained through the GEO2R online tool with |log FC| > 2 and adjust *p* value < 0.05. Then, the original data in TXT format is input into Venn online software to detect the common DEGs in two data sets. DEGs with log FC <0 are considered as down-regulated gene, while DEGs with log FC > 0 are considered as up-regulated gene.

### Gene Ontology (GO) and pathway enrichment (KEGG) analysis

Gene ontology analysis (go) is used to annotate and identify high-throughput transcriptome or genome data and analyze the biological process of these genes, and KEGG is used to analyze the biological function and pathway of DEG expression. Next we used David (annotation visualization and integrated discovery database, https://david.ncifcrf.gov/) (version 6.7) software to study the functional enrichment conditions of up-regulated and down regulated differential expression of DEGs, *p* <a 0.05 was considered statistically significant.

### PPI network and module data analysis

PPI information can be evaluated using the online tool STRING (earch tool for searching interacting genes, http://string-db.org) (version 11.0). Then, the STRING application in Cytoscape (version 3.7.2) was used to check the potential correlation between these DEGs (maximum interaction number = 0 and confidence score ≥ 0.4). Moreover, the plug-in MCODE application in Cytoscape was used to cluster the given network based on topology to find the area of dense connection^[7]^, which was used to identify the most important modules in PPI network (degree cutoff = 2, max. Depth = 100, k-core = 2, and node score cutoff = 0.2).

## Results

### Identification of DEGs in directly reprogrammed cardiomyocyte-like cells

In this study, there are 5 cardiomyocyte-like groups and 5 mouse cardiomyocyte groups. Through the GEO2R online tool, we extracted 2507 and 436 DEGs from GSE99814 and GSE22292, respectively. Then we use Venn diagram online software to make Venn diagrams of DEGs for dataset GSE99814 and GSE22292. The results showed that 243 DEGs were selected, including 127 up-regulated genes (log FC> 0) and 116 down-regulated genes (log FC< 0) between the iCMs and the mouse cardiomyocytes (table 1 & figure 1).

**Fig. 1.**
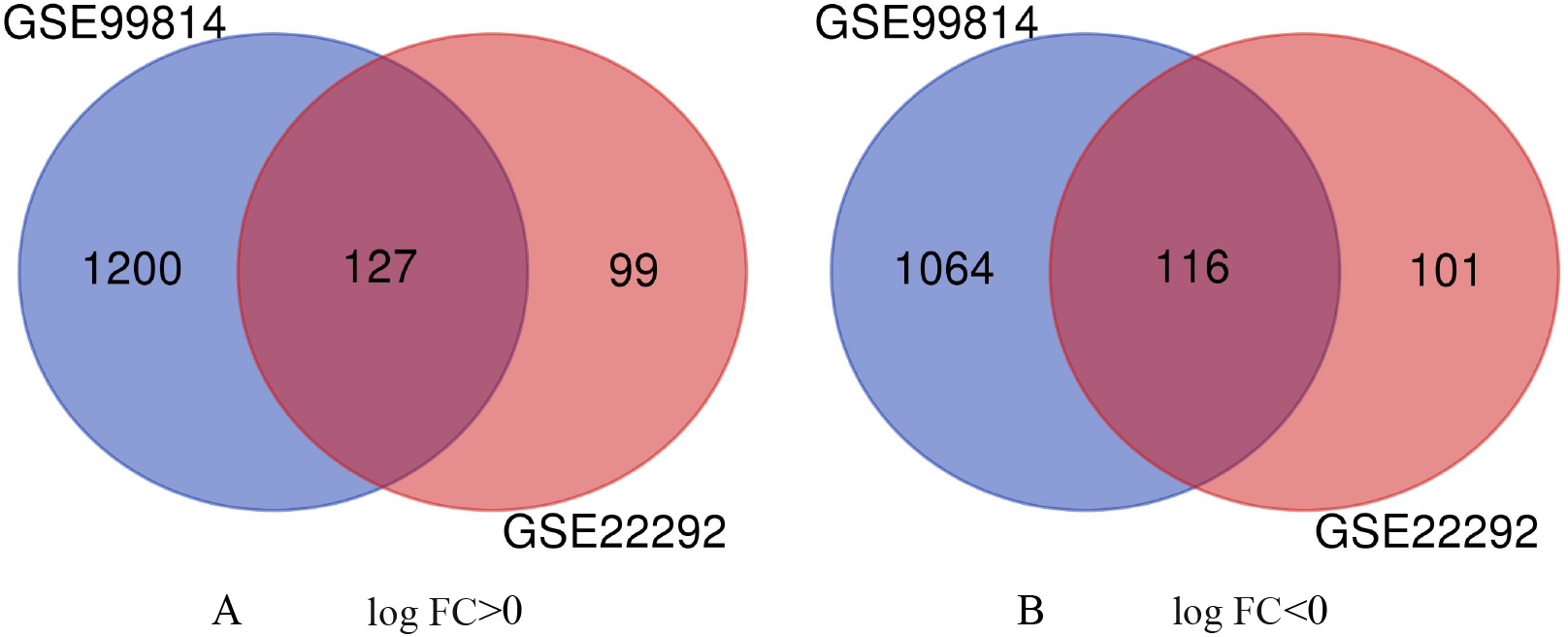
Through Venn diagram software (http://bioinformatics.psb.ugent.be/webtools/venn/), 243 common DEGs in two data sets (GSE99814 and GSE22292) were authenticated. Different colors indicated different datasets. A 127 protein expression was up-regulated (log FC > 0) in two datasets.. B 116 protein expression was down regulated (log FC < 0) in two datasets.

**Table 1.**
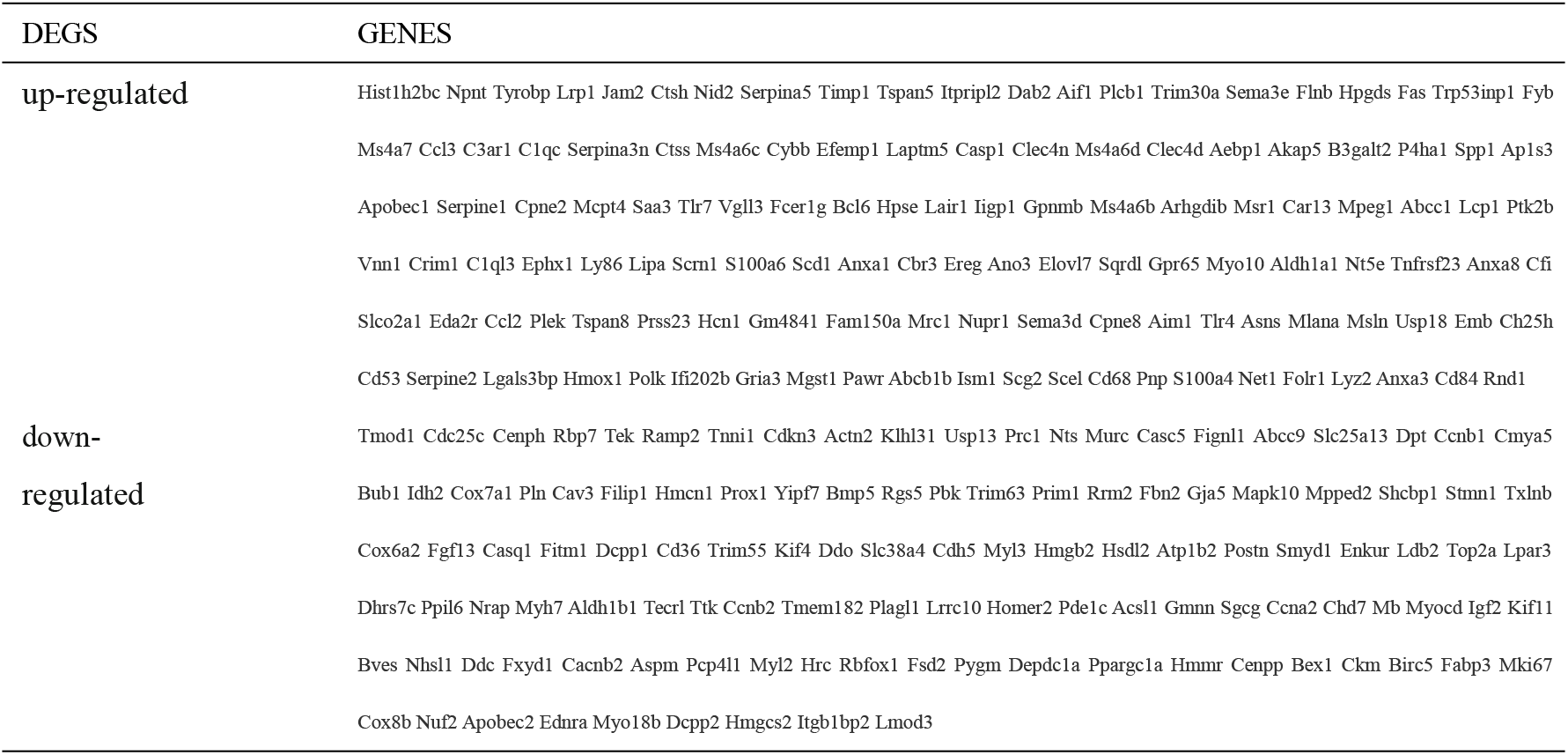
All 243 commonly differentially expressed genes (DEGs) were screened from two microarray datasets, including 127 up-regulated genes and 116 down-regulated genes.

### Gene ontology (GO) analysis of DEGs in directly reprogrammed cardiomyocyte-like cells

243 DEGs were analyzed by David software. The results of go analysis showed that 1) for biological processes (BP), the up-regulated DEGs were mainly concentrated in innate immune response, inflammatory response, immune system process, positive regulation of apoptotic process and cell adhesion, and down-regulated DEGs were particularly enriched in cell cycle, cell division, heart development, cardiac muscle contraction and cardiac muscle cell development; 2) for GO cell component (CC), the up-regulated DEGs were concentrated in membrane, plasma membrane, extracellular exosome, extracellular region and cytosol, while the down regulated DEGs were mainly concentrated in Z disc, cytoplasm, cytoskeleton, chromosome and sarcolemma; 3) for molecular function (MF), the up-regulated DEGs were enriched in protein binding, hydrolase activity, calcium binding, identical protein binding and actin binding, while the down regulated DEGs were in protein binding, nucleotide binding, calcium ion binding, actin binding and calcium channel regulatory activity (Table 2).

**Table 2.** Gene ontology analysis of differentially expressed genes in directly reprogrammed cardiomyocyte-like cells

| Expression | Category | Term | Count | % | p-Value | FDR |
| --- | --- | --- | --- | --- | --- | --- |
| Up-regulated | GOTERM_BP_DIRECT | GO:0045087~innate immune response | 16 | 0.0983 | 3.66E-08 | 5.82E-05 |
|  | GOTERM_BP_DIRECT | GO:0006954~inflammatory response | 15 | 0.0921 | 3.93E-08 | 6.24E-05 |
|  | GOTERM_BP_DIRECT | GO:0002376~immune system process | 12 | 0.0737 | 3.31E-05 | 5.26E-02 |
|  | GOTERM_BP_DIRECT | GO:0043065~positive regulation of apoptotic process | 11 | 0.0676 | 5.56E-05 | 8.82E-02 |
|  | GOTERM_BP_DIRECT | GO:0007155~cell adhesion | 11 | 0.0676 | 1.06E-03 | 1.6710 |
|  | GOTERM_CC_DIRECT | GO:0016020~membrane | 75 | 0.4607 | 3.75E-08 | 4.64E-05 |
|  | GOTERM_CC_DIRECT | GO:0005886~plasma membrane | 53 | 0.3256 | 2.00E-05 | 2.47E-02 |
|  | GOTERM_CC_DIRECT | GO:0070062~extracellular exosome | 42 | 0.2580 | 2.33E-08 | 2.89E-05 |
|  | GOTERM_CC_DIRECT | GO:0005576~extracellular region | 33 | 0.2027 | 2.60E-08 | 3.21E-05 |
|  | GOTERM_CC_DIRECT | GO:0005829~cytosol | 21 | 0.1290 | 8.06E-03 | 9.5337 |
|  | GOTERM_MF_DIRECT | GO:0005515~protein binding | 35 | 0.2150 | 3.02E-02 | 33.3590 |
|  | GOTERM_MF_DIRECT | GO:0016787~hydrolase activity | 18 | 0.1106 | 1.19E-02 | 14.6161 |
|  | GOTERM_MF_DIRECT | GO:0005509~calcium ion binding | 13 | 0.0799 | 1.19E-03 | 1.5609 |
|  | GOTERM_MF_DIRECT | GO:0042802~identical protein binding | 10 | 0.0614 | 1.47E-02 | 17.7513 |
|  | GOTERM_MF_DIRECT | GO:0003779~actin binding | 7 | 0.0430 | 1.79E-02 | 21.2252 |
| Down-regulated | GOTERM_BP_DIRECT | GO:0007049~cell cycle | 14 | 0.0767 | 3.32E-05 | 5.22E-02 |
|  | GOTERM_BP_DIRECT | GO:0051301~cell division | 11 | 0.0603 | 4.21E-05 | 6.62E-02 |
|  | GOTERM_BP_DIRECT | GO:0007507~heart development | 8 | 0.0438 | 6.16E-04 | 9.63E-01 |
|  | GOTERM_BP_DIRECT | GO:0060048~cardiac muscle contraction | 4 | 0.0219 | 2.37E-03 | 3.6597 |
|  | GOTERM_BP_DIRECT | GO:0055013~cardiac muscle cell development | 3 | 0.0164 | 3.03E-03 | 4.6620 |
|  | GOTERM_CC_DIRECT | GO:0030018~Z disc | 9 | 0.0493 | 3.59E-07 | 4.40E-04 |
|  | GOTERM_CC_DIRECT | GO:0005737~cytoplasm | 51 | 0.2794 | 2.77E-03 | 3.3391 |
|  | GOTERM_CC_DIRECT | GO:0005856~cytoskeleton | 14 | 0.0767 | 6.84E-03 | 8.0657 |
|  | GOTERM_CC_DIRECT | GO:0005694~chromosome | 8 | 0.0438 | 2.62E-03 | 3.1650 |
|  | GOTERM_CC_DIRECT | GO:0042383~sarcolemma | 7 | 0.0383 | 4.30E-05 | 5.27E-02 |
|  | GOTERM_MF_DIRECT | GO:0005515~protein binding | 38 | 0.2082 | 2.37E-03 | 3.0804 |
|  | GOTERM_MF_DIRECT | GO:0000166~nucleotide binding | 19 | 0.1041 | 3.11E-02 | 34.1068 |
|  | GOTERM_MF_DIRECT | GO:0005509~calcium ion binding | 10 | 0.0548 | 2.12E-02 | 24.6156 |
|  | GOTERM_MF_DIRECT | GO:0003779~actin binding | 7 | 0.0383 | 1.44E-02 | 17.4343 |
|  | GOTERM_MF_DIRECT | GO:0005246~calcium channel regulator activity | 3 | 0.0164 | 8.59E-03 | 10.7649 |

### Analysis of KEGG pathway in directly reprogrammed cardiomyocyte-like cells

Using David software to analyze 243 DEGs, and KEGG results (Table 3) showed that the up-regulated DEGs were significantly enriched in the four signaling pathways of Chagas disease (American trypanosomiasis), Lysosome, Complement and coagulation cascades and Salmonella infection, the down-regulated DEGs were enriched in the signal pathways of Cardiac muscle contraction, Cell cycle, Dilated cardiomyopathy, Adrenergic signaling in cardiomyocytes and Hypertrophic cardiomyopathy (HCM).

**Table 3.**
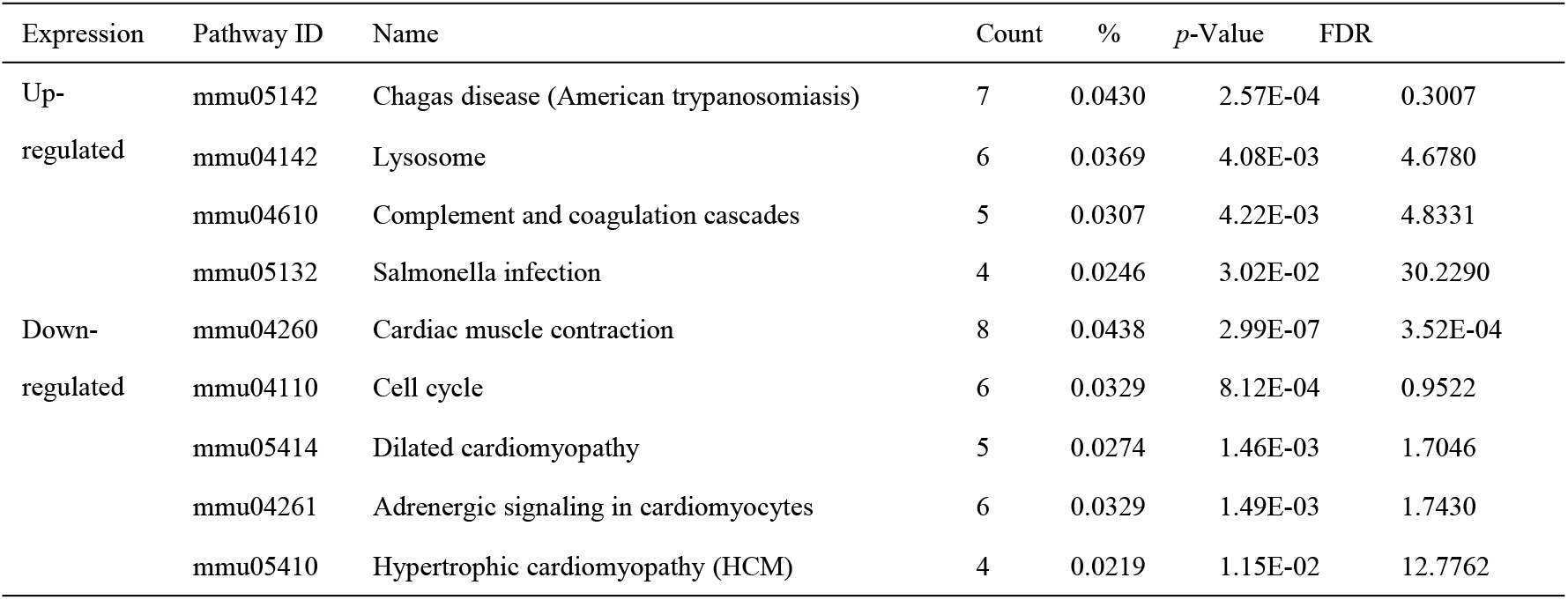
Signal pathway analysis of differentially expressed genes in directly reprogrammed cardiomyocyte-like cells

| Expression | Pathway ID | Name | Count | % | *p*-Value | FDR |
| --- | --- | --- | --- | --- | --- | --- |
| Up-regulated | mmu05142 | Chagas disease (American trypanosomiasis) | 7 | 0.0430 | 2.57E-04 | 0.3007 |
|  | mmu04142 | Lysosome | 6 | 0.0369 | 4.08E-03 | 4.6780 |
|  | mmu04610 | Complement and coagulation cascades | 5 | 0.0307 | 4.22E-03 | 4.8331 |
|  | mmu05132 | Salmonella infection | 4 | 0.0246 | 3.02E-02 | 30.2290 |
| Down-regulated | mmu04260 | Cardiac muscle contraction | 8 | 0.0438 | 2.99E-07 | 3.52E-04 |
|  | mmu04110 | Cell cycle | 6 | 0.0329 | 8.12E-04 | 0.9522 |
|  | mmu05414 | Dilated cardiomyopathy | 5 | 0.0274 | 1.46E-03 | 1.7046 |
|  | mmu04261 | Adrenergic signaling in cardiomyocytes | 6 | 0.0329 | 1.49E-03 | 1.7430 |
|  | mmu05410 | Hypertrophic cardiomyopathy (HCM) | 4 | 0.0219 | 1.15E-02 | 12.7762 |

### Protein–protein interaction network (PPI) and modular analysis

A total of the 243 DEGs, but 45 DEGs were not included in the PPI network and 198 DEGs were imported into the DEGs PPI network. Therefore, the PPI network were including 60 down-regulated genes and 44 up-regulated genes, which involving 198 nodes (genes) and 1173 edges. After, we used Cytoscape MCODE for further analysis. The results showed that there were 27 central nodes, all of which were down-regulated genes.

### Reanalysis of 27 MCODE genes by KEGG pathway enrichment

In order to understand the potential signal pathways of the 27 central genes, KEGG pathway enrichment of David software was used again to analyze the 27 genes (*p* < 0.05), among which 6 genes (CCNB1, CCNB2, BUB1, TTK, CDC25C, CCNA2) were enriched in the cell cycle pathway (*p*=1.20E-07, Table 4 & Fig. 3).

**Fig. 2.**
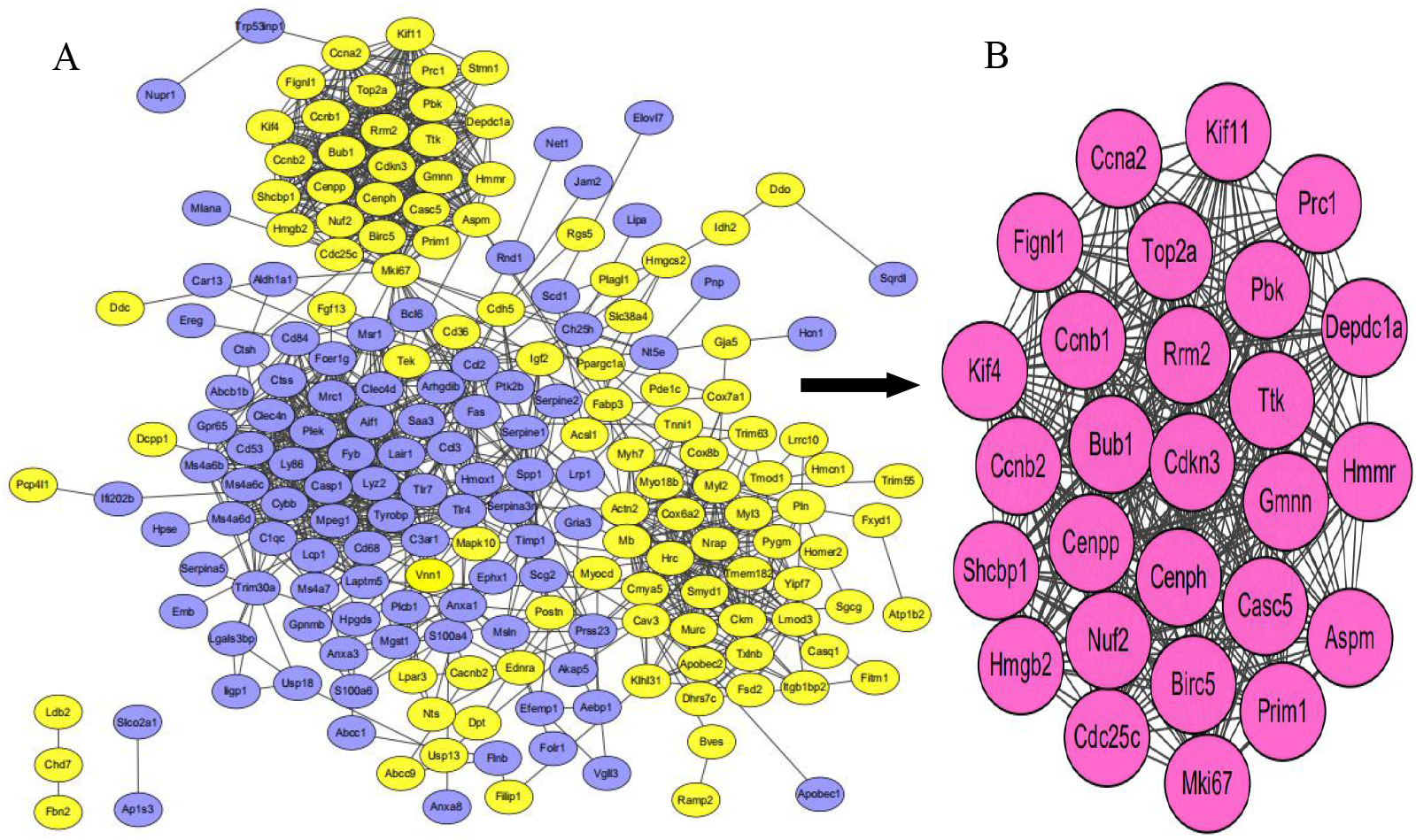
PPI network of common DEGs based on STRING online database and Module analysis.A There were 198 DEGs in the DEGs PPI network complex. The nodes represented proteins; the edges represented interaction of proteins; yellow circles represented down-regulated DEGs and blue circles represented up-regulated DEGs. B Module analysis through Cytoscape software (degree cutoff=2, nodes core cutoff=0.2, k-core=2, and max. Depth=100).

**Fig. 3.**
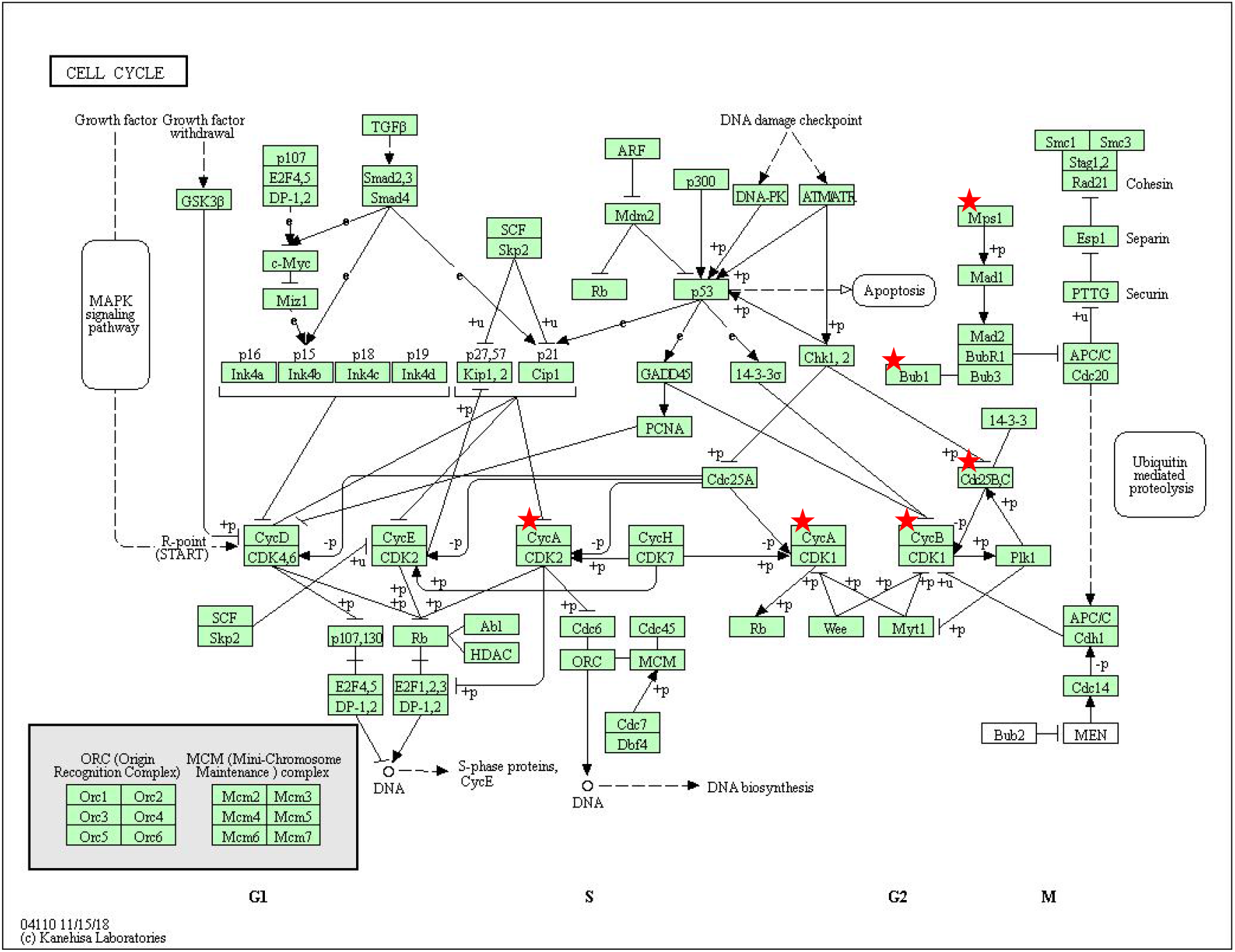
cell cycle signaling pathway. Six genes (CCNB1, CCNB2, BUB1, TTK, CDC25C, CCNA2, red stars) were significantly enriched in cell signaling pathway (*p*=1.20E-07).Mps1 means TTK, Cdc25B,C means CDC25C, CycB means CCNB1 and CCNB2, CycA means CCNA2.

**Table 4.** 27 genes were reanalyzed by KEGG pathway enrichment

| Pathway ID | Name | Count | % | p-Value | Genes |
| --- | --- | --- | --- | --- | --- |
| mmu04110 | Cell cycle | 6 | 0.1426 | 1.20E-07 | CCNB1, CCNB2, BUB1, TTK, CDC25C, CCNA2 |
| mmu04914 | Progesterone-mediated oocyte maturation | 5 | 0.1188 | 1.84E-06 | CCNB1, CCNB2, BUB1, CDC25C, CCNA2 |
| mmu04114 | Oocyte meiosis | 4 | 0.0951 | 2.37E-04 | CCNB1, CCNB2, BUB1, CDC25C |
| mmu04115 | p53 signaling pathway | 3 | 0.0713 | 2.59E-03 | CCNB1, CCNB2, RRM2 |

## Discussion

GSE22292 dataset from PubMed article PMID: <u>20691899</u>, Masaki et al.^[2]^ had demonstrated that the combination of three transcription factors GATA4, MEF2C and TBX5 can rapidly and effectively induce the newborn mouse cardiac fibroblasts into induced cardiomyocytes.

Interestingly, iCMs were electrophysiologically similar to neonatal mouse cardiomyocytes, and can contract spontaneously. And In the whole gene expression profile, it was also found that functional cardiomyocytes can be produced from differentiated somatic cells by certain factors. For example, the cardiac genes with important functions were significantly more up-regulated in 4-weekiCMs than in 2-week iCMs, but more up-regulated in neoCM than in 4-weekiCM or cardiac fibroblasts. Therefore, in terms of gene expression profile, iCMs were similar to but different from cardiomyocytes of newborn mice. From the level of q-PCR, lysine 27 (H3K27me3) decreased significantly in the promoter after reprogramming, while lysine 4 (H3K4me3) increased in the promoter region of iCMs, and they were reaching the same level as that of cardiac cells. Generally, the chromatin state of iCM derived from cardiac fibroblasts is similar to normal cardiac cells in some cardiac specific genes.

GSE99814 data set from article PMID: 28954220, Zhou et al.^[8]^ compared and analyzed the molecular characteristics of reprogrammed contraction CM produced by iPSC differentiation or direct reprogramming of cardiac fibroblasts (CF) from the same source. The detection of fibroblast genes showed that H3K4me3 in iCM was higher than in iPSC-CM.Moreover, they evaluated the maturity of reprogrammed cm. GSEA analysis of neoCM and iCM showed that compared with the genome of adult CM, iCM had a higher correlation with neoCM. With the progress of iCM reprogramming, the score of gene set enrichment of CM and early embryo related genes decreased, but the score of gene set enrichment of mature CM related genes increased. This could mean that rapid maturation may occur during iCM induction, and the directly reprogrammed iCM seems to acquire adult-like CM function at the transcription level.

In this study, two gene expression databases (GSE99814 and GSE22292) of iCM induced by CF from the same source were analyzed by bioinformatics, and five groups of direct reprogramming iCMs and five groups of normal mouse cardiomyocytes were studied to identify the effective biomarkers for conversion during direct reprogramming. Through GEO2R and Venn software, we found 243 changed DEGs (| log FC |> 2 and adjusted p value < 0.05), and including 127 up-regulated DEGs (log FC > 0) and 106 were down regulated DEGs (log FC < 0). Then we used David to analyze the concentration of GO and KEGG enrichment analysis in 243 DEGs: 1) for biological processes (BP), the up-regulated DEGs were mainly concentrated in innate immune response, inflammatory response, immune system process, positive regulation of apoptotic process and cell adhesion, and down-regulated DEGs were particularly enriched in cell cycle, cell division, heart development, cardiac muscle contraction and cardiac muscle cell development; 2) for GO cell component (CC), the up-regulated DEGs were concentrated in membrane, plasma membrane, extracellular exosome, extracellular region and cytosol, while the down regulated DEGs were mainly concentrated in Z disc, cytoplasm, cytoskeleton, chromosome and sarcolemma; 3) for molecular function (MF), the up-regulated DEGs were enriched in protein binding, hydrolase activity, calcium binding, identical protein binding and actin binding, while the down regulated DEGs were in protein binding, nucleotide binding, calcium ion binding, actin binding and calcium channel regulatory activity; 4) the up-regulated DEGs were significantly enriched in the four signaling pathways of Chagas disease (American trypanosomiasis), Lysosome, Complement and coagulation cascades and Salmonella infection, and the down-regulated DEGs were enriched in the signal pathways of Cardiac muscle contraction, Cell cycle, Dilated cardiomyopathy, Adrenergic signaling in cardiomyocytes and Hypertrophic cardiomyopathy (HCM).

In order to further study the function of the identified DEGs, we used STRING and Cytoscape to build a PPI network with 198 nodes and 1173 edges. Then 27 core genes (Ttk, Top2a, Shcbp1, Rrm2, Prim1, Pr1c, Pbk, Nuf2, Mki67, Kif4, Kif11, Hmmr, Hmgb2, Gmnn, Fignl1, Depdc1a, Cenpp, Cenph, Cdkn3, Cdc25c, Ccnb2, Ccnb1, Ccna2, Casc5, Bub1, Birc5 和 Aspm) were screened from PPI network by MCODE analysis. At last, we used David software to analyze the KEGG pathway of the 27 important genes again, and found that 7 genes were enriched in the cardiac contraction signal pathway, and 6 genes were significantly enriched in the cell cycle pathway. These genes and signal pathways may be the key genes to induce CF into iCM through direct reprogramming.

Adult mammalian cardiomyocytes (CM) have a limited proliferative capacity, which makes the heart very inefficient in the formation of new cardiomyocytes when contractile cardiomyocytes are lost (such as myocardial infarction, dilated cardiomyopathy and other diseases) ^[9]^. After birth, cardiomyocytes continue to proliferate in a short neonatal period, which is the last peak of heart growth and the key to regeneration of damaged heart in newborn mice ^[10]^. Then, most cardiomyocytes withdraw from cell cycle and stop proliferation after puberty (7-10 years old) ^[11]^. However, amphibians have certain regeneration ability. For example, the larva of Xenopus Africana shows a decreased regeneration ability with age, in the early stage (50-53) before metamorphosis, amputation in the developing limb can also make the limb regenerate perfectly, but in the later stage of development, with the development of the immune system, it has a certain inhibitory effect on the regeneration ability. If amputation in this period, it will gradually lead to incomplete tail or limb development^[12]^. In 2015, Han et al.^[10]^ found that acute inflammatory response was induced by cardiac injury after apicectomy (AR) in newborn mice. Acute inflammation can induce the proliferation of cardiomyocytes in the heart of newborn mice. However, after immunosuppression, the proliferation of cardiomyocytes in AR was inhibited. And in the absence of interleukin 6 (IL-6), it can cause neonatal mouse cardiac injury to fail to reactivate and proliferate. The results showed that acute myocarditis was necessary to stimulate the proliferation of cardiomyocytes in newborn mice and enough to stimulate the proliferation of cardiomyocytes. This is consistent with the results of the DEGs upregulated by the CF direct reprogramming process in this study mainly enriched in the innate immune response, inflammatory response, and immune system process. So in mammals, immunological disorders may be the key to limiting the potential for regeneration.

In 2016, Zhang et al. ^[13]^ in order to determine the differential gene expression during regeneration, and they analyzed the cardiac transcripts of the first and seventh day after removal of the apex of the heart in newborn mice through RNA-seq. The results showed that on the first day, the genes involved in inflammatory response and wound healing were up-regulated, and also reflecting the acute infiltration of immune cells. On the contrary, cell cycle genes were activated, while sarcomere and cardiac contraction related genes showed lower expression at 7th day. The result of our study is that the genes involved in cell cycle have been down regulated, which may be that the sequenced transcriptome was reprogrammed and cultured for 4 weeks and developed into mature cardiomyocytes, and develops to the state of mature cardiomyocytes. At this time, it may have exited the cell cycle ^[14]^. Moreover, CFs can be induced into iCMs ^[2–4]^ with spontaneous shrinkage and beat in different reprogramming methods.

Therefore, abounding evidences have proved that the gene expression profiles of iCMs induced by direct reprogramming and CMs are very similar, and especially play an important role in cardiac reprogramming in the immune related genes, myocardial contraction and cell cycle signaling pathway. Perhaps in the near future, it is possible to induce CF into perfect normal cardiomyocytes by direct reprogramming with overcoming immunologic barriers and other genetic characterization.

## Conclusions

In this study, we compared the data of five groups of directly reprogrammed cardiomyocytes and five groups of normal mouse cardiomyocytes in two different microarray databases by bioinformatics. Through the identification of the DEGs, we found that the up-regulated genes are closely enriched in the immune related biological process, while the down regulated genes are mainly concentrated in the cell cycle signal pathway. The results show that they could play an important role in direct cardiac reprogramming. Sincerely, it is hoped that our research can provide some new directions and ideas for further improvement of direct cardiac reprogramming that can generate iCMs and cardiac regeneration medicine.

## Footnotes

Liaoning provincial key R & D project (2018225013); Shenyang Medical College Excellent Talents project (20113067); General project of National Natural Science Founding (81070128); Shenyang young and middle-aged innovation support project (RC180379);Graduate student innovation funding project ofShen Yang Medical College (Y20190501, Y20190507)

